# Sleep increases propagation speed of physiological brain pulsations

**DOI:** 10.1101/2025.01.09.632145

**Authors:** Ahmed Elabasy, Heta Helakari, Tommi Väyrynen, Zalán Rajna, Niko Huotari, Lauri Raitamaa, Ville Isokoski, Matti Järvelä, Mika Kaakinen, Johanna Piispala, Mika Kallio, Vesa Korhonen, Tapio Seppänen, Vesa Kiviniemi

## Abstract

During sleep, there is an increase in the brain cerebrospinal fluid (CSF) solute convection driven by physiological pulsations. Although the main drivers of CSF flow, namely cardiac, respiratory, and vasomotor pulsations, become more powerful during sleep, there is relatively little information regarding their effects on CSF flow velocity across human brain during sleep. Here, we used functional magnetic resonance encephalography (MREG) to measure non-invasively changes in brain water flow during to sleep by tracking the propagating ultrafast signal changes induced by physiological brain pulsations. We first undertook a phantom study confirming that dense optical flow analysis of MREG data accurately detects water flow velocity, and reflects the power of the physiological pulsations. We then applied the method to quantify CSF water flow velocity in brain of healthy volunteers during EEG-verified awake and sleep recordings of ultrafast MREG data. Sleep induced an increase in CSF flow speed, as demonstrated by elevated vasomotor and respiratory pulsation speeds, while the speed of cardiovascular impulse propagation remained unchanged. The speed increases match previous findings of respective pulsation power changes, and correlated with slow delta EEG power. The sleep-induced CSF flow speed increases occurred dynamically over both pulsation cycles, without the large effects on flow directions reported previously in several neurological conditions. In conclusion, sleep increases 3D water flow speed dynamically in human brain regions showing concomitant pulse power increases, supporting a porous media model of hydrodynamics in brain cortex.

## Background

Recent investigations have shown that glymphatic mechanisms increase interstitial and perivascular brain solute convection during sleep. Tracking the distribution in rodent brain of fluorescent tracers by multi-photon microscopy or gadolinium tracers by MRI *in vivo*, following their injection into CSF spaces reveals the cortical glymphatic solute convection (1,2). Applying such contrast media enables the imaging of glymphatic solute convection in brain of experimental animal and human volunteers (3–7). These methods depict the overall tracer kinetics extending over an interval of several minutes to hours, based on the net effects of the physiological drivers of the glymphatic solute convection within the CNS. Due to the high frequency of physiological drivers with respect to slow movement of tracers, contrast methods cannot reveal dynamic changes in the forces driving CSF solute convection in the human brain.

Physiological pulsations propagating along blood vessels and cerebrospinal fluid (CSF) spaces are the drivers of glymphatic solute convection (4). In living mice, cardiovascular pulsations drive solute transport in para-arterial CSF space and sub-arachnoid spaces (3,4), while respiratory pulsations predominantly affect (peri)venous pulsations (8–11). Very low-frequency (VLF) vasomotor waves of arterial smooth muscle have recently been suggested to also convect solutes (9,12,13).

Ultrafast functional MRI (fMRI), utilizing magnetic resonance encephalography (MREG) sequences, enables non-invasive tracking of physiological impulses in the brain by following magnetic spin perturbations in water protons. This technique can simultaneously quantify the power, speed, and direction of these impulses as they propel water molecules through CSF spaces and blood vessels (9, 14–17). The physiological pulsations can be critically sampled with the ultrafast 10 Hz (i.e., 10 whole 3D brain scans per second) sampling of T2* weighted brain water signal, which enables the accurate simultaneous acquisition of cardiovascular pulsation ≈ 1 Hz, respiratory pulsation ≈ 0.3 Hz, and vasomotor driven very low frequency (VLF) pulsations (<0.1 Hz) without aliased mixing of the three pulsations. This is in contrast to classical fMRI, where the physiological pulsations alias over the low frequency range of BOLD signal due to the non-critically low image repetition times TR > 500 ms (9, 18, 19).

During human sleep, the powers of all physiological brain pulsations increase as a function of sleep depth in brain regions of slow delta activity, this in conjunction with increased CSF flushing (20,21). Other research indicates that obesity, as measured by body mass index (BMI), has significant associations with physiological brain pulsations, with notably increased respiratory pulsations, along with decreased pulsations related to arterial function in key brain structures such as the pituitary gland and hypothalamus (22). In addition, multiple brain diseases such as narcolepsy, epilepsy, and Alzheimer’s disease manifest unique alterations in the power of physiological brain pulsations (16, 17, 23–25).

From the perspective of physics, the power (or amplitude) of a wave has no influence on the propagation speed in free-flowing liquid. However, in a porous medium like the brain, which contains innumerable (peri)vascular fluid conduits flowing in and out of the brain tissue, the power of a wave does influence the propagation speed (**v_s_**) (26–28). Optical flow analysis of MREG data (15,17) was able to quantify several forms of changes in both the speed and direction of the cardiovascular pulsations in Alzheimer’s disease that anatomically follow the distribution of each arterial territory (16). A recent study showed slowing of the propagation of respiratory impulses in brain of patients with epilepsy (17). In addition to altered flow velocity magnitude (**v_s_**), also the directional (<u>v</u>) trajectories of impulse propagation show differences in neuropathological conditions (16,17).

In this study, we first verified in a phantom experiment that MREG indeed accurately depicts the pulsation speed of water flow, and that the optical flow analysis MREG data thus accurately quantifies water molecule flow. We then recorded for the first time the propagation velocity (V, i.e. both **v_s_** and <u>v</u>) of all three physiological brain pulsations in a group of healthy volunteers. Given the known associations between sleep stage and pulsation power, we next investigated the effect of sleep on pulsation V in the sleeping human brain. We thus tested the hypothesis that the direction and speed of brain tissue pulsations would change during sleep, in relation to increased solute convention. To this end, we used ultrafast MREG imaging to quantify the changing magnitude (**v_s_**) and directional (<u>v</u>) trajectories in CSF and brain tissue for each physiological brain pulsation during awake vs. EEG-verified NREM sleep state. Results show compelling evidence of how physiological water flow pulsations speed up in natural human sleep.

## Materials and methods

### Subjects

The regional medical research ethics committee of the Wellbeing services county of North Ostrobothnia approved the study. All participants gave written informed consent, according to the requirements of the Declaration of Helsinki. We first interviewed and screened prospective participants. Subjects reported good general health, and met the following inclusion criteria: non-smokers without neurological or cardio-respiratory disease, no continuous medication, and no possibility for pregnancy. Subjects did not have contraindications or metal in the body that would have precluded MRI scanning.

We scanned a group of 22 subjects (27.2 ± 4.9 years, 12 females) during an awake control state in the afternoon (4-6 PM) after a normal night’s sleep. Twelve subjects (26.2 ± 4.3 years, 5 females) were scanned while asleep (6-8 AM) after a full night of sleep deprivation, and ten subjects (28.4 ± 5.7 years, 7 females) while asleep during normal sleep time (10-12 PM). We recorded sleep scoring in all participants. As the VLF < 0.1 Hz wave speed analysis requires data extending over several wave cycles, we used the 5-minute data epoch with the highest amount of sleep. We present the pipeline for analyzing the pulsation propagation velocity of study subjects in a study flow chart (Figure 1).

**Figure 1.**
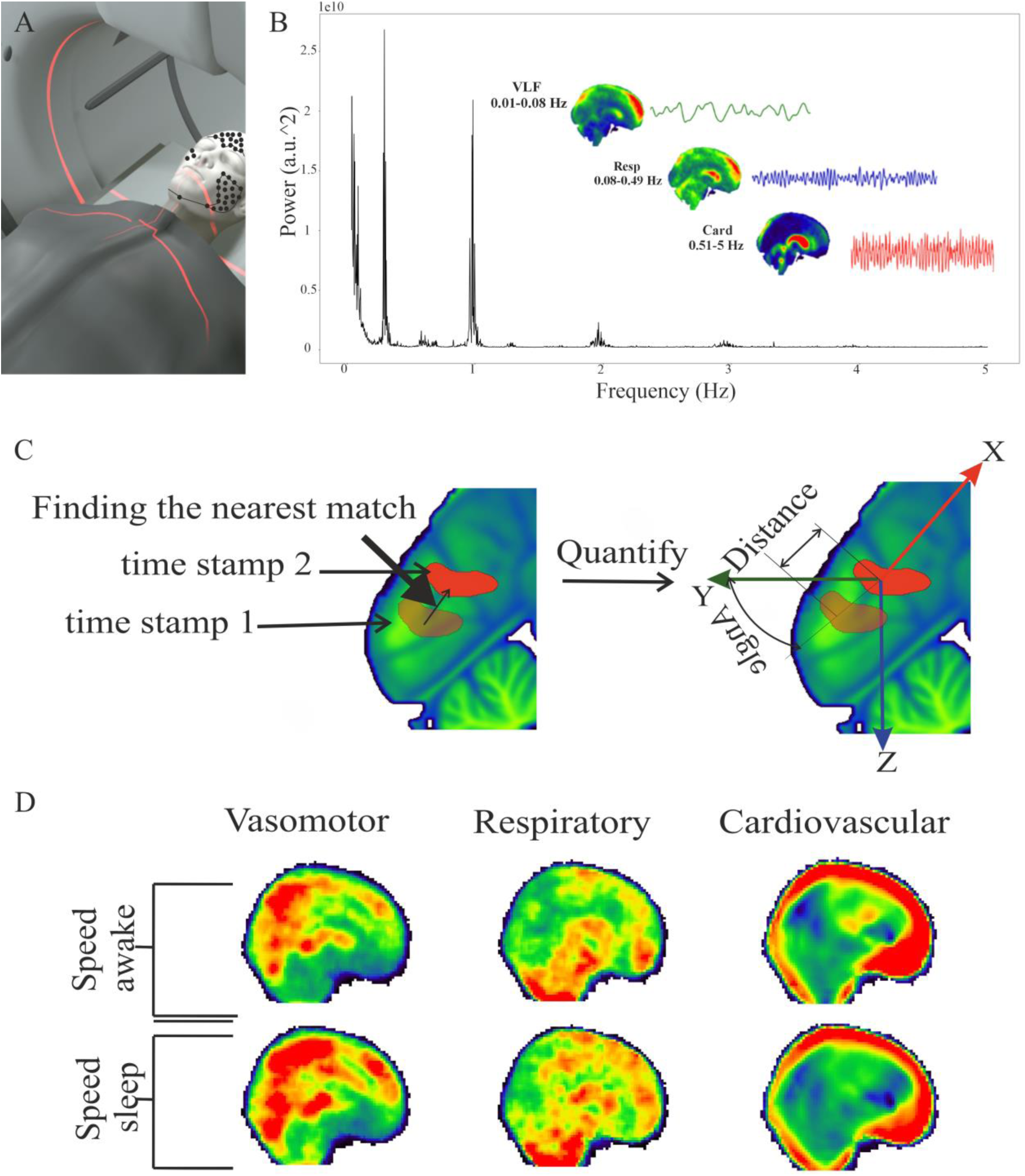
Flow chart for velocity analysis of study subjects. (A) Simultaneous multimodal MREG recordings with a 3T fMRI and dcEEG (EGI ch 256) data collection in awake and sleep states on separate days. (B) FFT power spectral analysis illustrating the distinct signal-to-noise characteristics of the vasomotor, respiratory, and cardiovascular brain pulsations. The images illustrate power distributions of these pulsations in MNI152 space. Each pulsation is illustrated on the right after an individual band-pass filtering of the MREG data to separate the cardiac, respiratory, and vasomotor signals in their own frequency bands. (C) 3D depiction of optical flow analysis of band-passed data following signal changes between consecutive MREG images obtained 100 ms apart. (D) Visualization of speed results in MNI space.

### Data acquisition

All subjects were scanned in Oulu University Hospital (Finland) using a Siemens MAGNETOM Skyra 3T scanner (Siemens Healthineers AG, Erlangen, Germany) equipped with a 32-channel head coil. We applied the fast fMRI sequence known as MREG (29–32) in synchrony with a previously described multimodal scanning setup (33). MREG is a single-shot, critically sampled (10 Hz sampling rate) sequence that allows visualizing the cardiovascular (≈ 1 Hz), respiratory (≈ 0.3 Hz), and very low frequency (VLF) vasomotor pulsations (<0.1 Hz), without higher frequencies to alias on the top of the lower frequencies (9).

The following parameters were used for MREG: repetition time (TR) = 100 ms, echo time (TE) = 36 ms, and field of view (FOV) = 192 mm, 3 mm cubic voxels and flip angle (FA) = 5°. Furthermore, the magnetization crusher gradient between scans was set to 0.1, to avoid masking of the signal by stimulated slow echo drifts, while retaining good sensitivity to physiological pulsations. Parameters for three-dimensional structural T1 MPRAGE were: TR = 1900 ms, TE = 2.49 ms, FA = 9°, FOV = 240 mm, and slice thickness 0.9 mm. We obtained a structural T1 MRI scan in all sessions, for registering the fMRI data to the standard MNI152 3 mm brain template for group analyses.

The Electrical Geodesics (EGI, Magstim Company Ltd, Whitland, UK) MR-compatible GES 400 system, equipped with a 256-channel high-density net, was used to record EEG in synchrony with MREG at settings as previously described (33). Respiratory belt and fingertip peripheral (SpO_2_) signals from the Skyra scanner, along with, ECG, SpO2, and end-tidal carbon dioxide (EtCO2) from an anesthesia monitor (Datex-Ohmeda S/5 Collect software) were measured in synchrony with the MREG/EEG recordings (33).

### Preprocessing and analysis of EEG data

EEG data were preprocessed using the Brain Vision Analyzer (Version 2.1; Brain Products) as described in (20). After correction of the MRI gradient and for ballisto-cardiographic artifacts, two experienced clinical neurophysiologists (JP, MK) performed consensus sleep scoring according to the 10-20 system in 30 s epochs, following AASM guidelines for clinical sleep studies (American Academy of Sleep Medicine, 2017). Their viewing of 10-20 minutes of sleep data from each subject mainly scored as N1 or N2 stage sleep. They also confirmed wakefulness by scoring the awake data, which indicated a single epoch of drowsy N1 sleep in the awake dataset; wakefulness data was missing from two subjects. The present sleep scoring method is consistent with that applied in previous research (20, 21, 34).

For the optical flow analysis, we selected a 5-minute segment with fully awake, and another 5-minute segment with the highest amount of sleep, rating as N1-N2, depending on the individual subject.

As we have shown earlier that the MREG pulsation power increases spatially in the same areas as the slow-delta increases (20) and that there is a general linear relationship between these powers (35), we wanted to investigate whether the slow-delta power is related also to the speed of the pulse propagation in addition to power. For studying the relation between MREG power and slow-delta power from the EEG data, we first filtered the EEG data to extract the slow-delta power (0.2-2 Hz), which supports an evaluation of sleep quality. To this end, we computed the mean the total slow-delta power for all subjects, after excluding the noisy EEG channels using all methods implemented in PyPREP Version 0.4.2 (36), mainly consisting of a random sample consensus approach (RANSAC) to identify bad EEG channels (37). We excluded the EEG data of one subject due to corrupted signal. We computed the log of the MREG power sum versus the log of the slow-delta power sum and filtered out all outliers using the strict four standard-deviation acceptance threshold according to Chebyshev’s theorem.

### Preprocessing of MREG data

We undertook image reconstruction using a L2-Tikhonov regularization with lambda value 0.1, where we had determined the latter regularization parameter by the L-curve method with a MATLAB recon-tool provided by the sequence developers (38). Before preprocessing, the image reconstruction also included a critical dynamic off-resonance in k-space (DORK) step, which corrected for scanner warming and respiration-induced dynamic B_0_-field changes (39,40).

After reconstruction, we preprocessed and analyzed the MREG data using the Functional MRI of the Brain Software Library (FSL; Brain Extraction Tool, version 5.09) (41), and Analysis of Functional NeuroImages (AFNI, version 2) (42) in MATLAB (R2021) and Python 3.7.3. We first extracted the brain from structural 3D MPRAGE volumes using neck cleanup and bias field correction options (41). We then used the FSL pipeline to preprocess the functional data. The pipeline included high-pass filtering with a cutoff frequency of 0.008 Hz, spatial smoothing with 5 mm full width at half-maximum (FWHM) Gaussian kernel, and motion correction (FSL 5.08 MCFLIRT) (43). We found no significant differences between awake and sleep state in the relative or absolute mean displacement values (mm) (p_all_ > 0.094). Nonetheless, we followed the motion correction by applying the 3dDespike function in AFNI (Analysis of Functional NeuroImages, v242) to remove the highest spikes in the MREG data time series. We then registered the preprocessed data to the MNI152 space at 3-mm resolution, to allow comparable analysis between subjects.

We had obtained 10 minutes of awake data and 10-20 minutes of sleep data from all subjects. For the final analysis, we used fslroi to segment the data into the best 5-minute awake and sleep epochs, according to EEG. For most of the subjects, we used the first 5 minutes of awake data to confirm wakefulness. For one subject, we chose the last 5 minutes, because the first 5 minutes included some sleep, and for another subject we were obliged to use awake data containing an epoch of EEG confirmed sleep. For sleep analysis, we chose a 5-minute epoch that included the highest amount of EEG-scored sleep (N1 and N2 sleep), as presented in Results.

### Respiratory and cardiovascular pulsation ranges

To calculate optical flow in physiological pulsations, we verified with MATLAB (R2021b) the individual respiratory and cardiac frequency ranges from physiological EtCO_2_ (or respiratory MR scanner belt signals) and fingertip SpO_2_ signals from the FFT spectra of physiological respiratory and cardiac signals. We chose the smallest minimum and highest maximum detected respiratory rate values over the whole group (0.08-0.49 Hz), c.f. Table 1. Cardiovascular frequency was high-passed from 0.51 to the full 5 Hz range for the whole group, in order to capture harmonics that represent a realistic cardiovascular impulse shape, as required for precise velocity detection (16). For very low frequency, we used the range (0.01-0.08 Hz). We chose the minimum value based on the typical threshold values reported in the literature, and based the maximum value on the lowest respiratory values present in our data, thereby minimizing the overlapping effects of respiration and vasomotion.

**Table 1.**
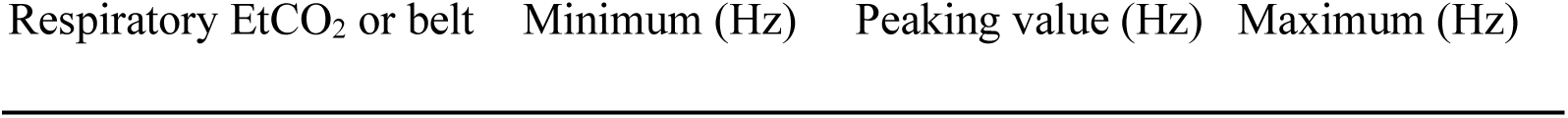
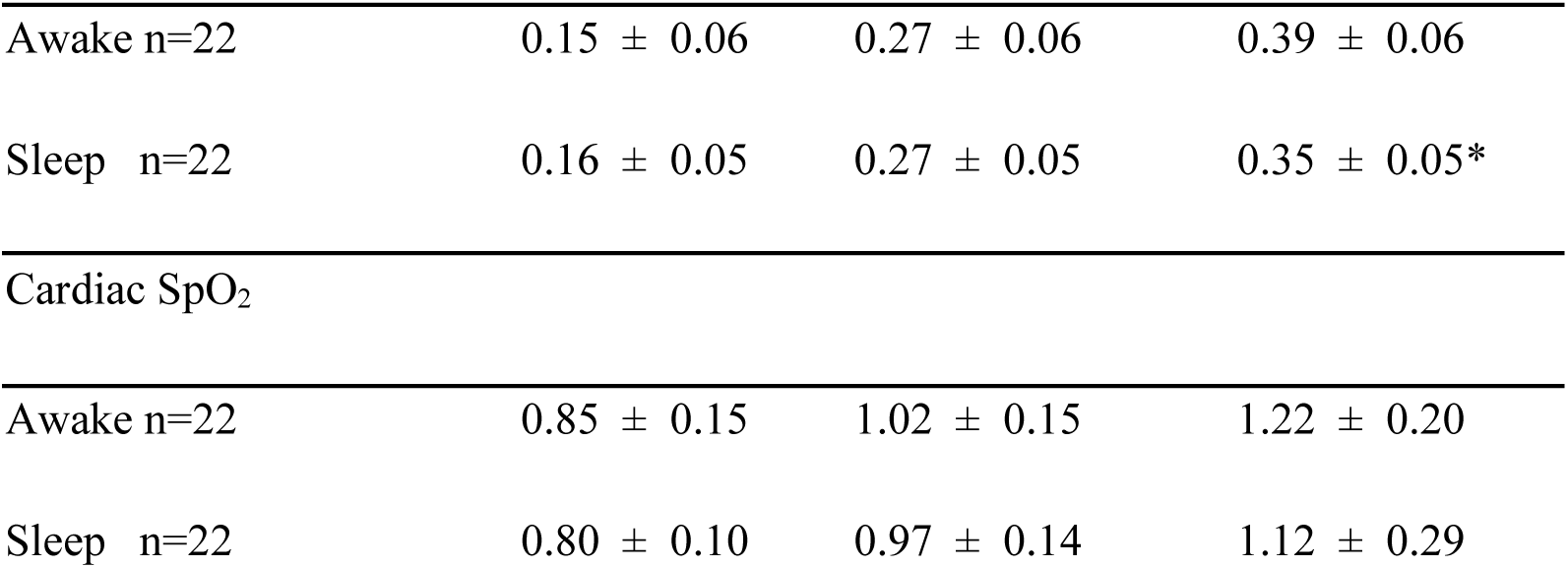
Respiratory and cardiac frequency ranges as determined from physiological EtCO2 or respiratory belt and SpO2 signals. Results are expressed as mean ± standard deviation for the minimum, peaking and maximum values in (n=22) recordings. Statistical differences between awake and sleep states are expressed with * p < 0.05.

| Respiratory EtCO<sub>2</sub> or belt | Minimum (Hz) | Peaking value (Hz) | Maximum (Hz) |
| --- | --- | --- | --- |
| Awake n=22 | 0.15 ± 0.06 | 0.27 ± 0.06 | 0.39 ± 0.06 |
| Sleep n=22 | 0.16 ± 0.05 | 0.27 ± 0.05 | 0.35 ± 0.05* |
| Cardiac SpO<sub>2</sub> |  |  |  |
| Awake n=22 | 0.85 ± 0.15 | 1.02 ± 0.15 | 1.22 ± 0.20 |
| Sleep n=22 | 0.80 ± 0.10 | 0.97 ± 0.14 | 1.12 ± 0.29 |

### Optical flow analysis

The Lucas-Kanade multi-resolution 3D optical flow method has proved to be an efficacious for tracking cardiac impulses within the brain (15). Separately, we can thus quantify the following parameters: 3D direction <u>v</u> and speed **v_s_**, as well as their combined value, i.e. the total velocity V (16). Specifically, as can compute V as:

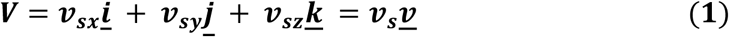

where 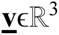, and **<u>i</u>, <u>j</u>, <u>k</u>** are the three standard unit vectors in the principal X, Y, and Z directions, respectively.

The magnitude of V represents the velocity vector in 3D. Every vector in 3D can be represented by the sum of the dot product of its magnitude in the X-direction multiplied by a unit vector (a vector with magnitude of one cm/s), in the X-direction, the dot product of its magnitude in Y-direction multiplied by a unit vector in Y-direction, and the dot product of its magnitude in Z-direction multiplied by a unit vector in Z-direction. Thus V can be represented by the dot product of the magnitude **v_s_,** which equals 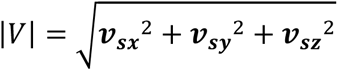, and the unit direction 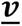,which equals 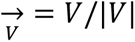. In the results section, the V map illustrates speed as the vector length and its direction, while the **v_s_** map illustrates only the combined speed in 3D.

Each cardiac brain impulse induces a clear MREG signal dip (from momentarily desynchronized spins) that propagates into the CSF and tissue from the cerebral arteries, most distinctly from major cerebral arteries including the anterior cerebral artery (ACA) (9, 14, 16, 18). This moving cardiac brain pressure wave is trackable with steps of sparse optical flow analysis, as in (16). While our previous approach was effective in tracking the cardiac signal (cardiac rate about 80 bpm, i.e. 1.3 Hz), it entailed considerable data loss due to the restricted use of 0.9 s cardiac cycles only. However, this type of restriction of slower periods like respiratory especially VLF induces unacceptable loss of information, because of the greater variability of the slower pulsations due to physiological signal variability. Our previous study overcame this problem by resampling all the velocity profiles of wave fronts extracted from the slower respiratory data (17).

The three sources of physiological cerebrovascular pulsations are cardiovascular impulses, vasomotor waves from all cerebral arteries, and respiratory pulses from opposed phase venous blood vs. CSF space pulsations inside the cranium (44). As liquids do not compress, the impulses immediately affect the (peri)vascular CSF water molecules, which then experience phase shifts at different speeds, depending on the frequency and magnitude of the physiological pulses. While sparse optical flow follows a single moving feature in the data, like the cardiovascular impulse nadir, it has limited capacity to capture displacement features. Here, we resort to dense optical flow analysis, which can track *all* signal changes between image frames as movement in the human MREG data, and thus captures much finer features of the 3D water movement (45, c.f. nanonets.com). In the time domain, dense optical analysis follows all changes over the entire wave pattern, rather than the separately detected signal minima observed in the sparse analysis. In the present study, by using pulsatile water-flow phantom measurements, we first show that dense optical flow analysis of MREG data is accurate in flow speed measurements and then present dense optical flow obtained with resampling of all three physiological human brain pulsations.

Given this background, to obtain a more comprehensive summation of the 3D movement of brain pulsations, we used the dense optical flow analysis of MREG data for the cardiovascular, respiratory, and vasomotor frequencies. Then, we resampled the optical flow results of cardiac pulses to 0.9 s and the respiratory pulsations to 6 s, based on our previous studies (16,17). For vasomotor waves, we used a 20 s wave cycle length, based on a pioneering study of the propagation of vasomotor BOLD waves (46).

For pulse triggering of the three physiological pulsations, we used the intracranial regions of interest with the most conspicuous and powerful pulsations, i.e., the anterior cingulate area (ACA) for cardiovascular (16), the IV ventricle for respiratory (17), and the posterior cingulate cortex (PCC) for vasomotor waves (46). Appropriate multi-resolution pyramid depth was used to compute the propagation speed, with scales of 3 for cardiac, 1 for respiratory, and zero for the VLF pulsations. This configuration enabled us to capture the gradual propagation of brain BOLD signals. For comparison, we also analyzed the fastest cardiovascular pulsations using both sparse and dense optical flow methods.

### Statistical analysis of velocity

We calculated a mean optical flow velocity magnitude (**v_s_**) map for each subject in 3D MNI space over the 0.9, 6 and 20 s cycles, respectively. We then performed a paired statistical analysis of the **v_s_** maps between awake and sleep states using FSL-TFCE (Threshold-Free Cluster Enhancement) randomise (threshold p < 0.05, n=5000) test sequentially for every time frame of the cycles, i.e. 9, 60 and 200 3D volumes. This yielded statistical maps for, respectively, v**_s_**_card_, v**_s_**_resp_ and v**_s_**_vaso_ over the cycles to provide a dynamic view of the pulsation speed changes, as in our previous work (17).

We likewise analyzed the directional velocity <u>v</u> maps analysis to find the reverse in flow directions in the contrast of sleep and awake states of the subjects. Here, we separated the X-, Y-, and Z-components of the vectors, and compared separately the positive and negative propagation directions of every component. We combined the resultant maps to depict the reversed <u>v</u> between the two states of arousal states as in (17). We depicted the results in 4 segments within the cycle, overlaid upon 1 mm MNI152 space.

### Power analysis

We utilized the 3dPeriodogram function in AFNI to compute an FFT power density map for each MREG dataset, separately for each individual and condition. The total number of time points across all scanning segments was 2861, and we conducted the FFT analysis with 4096 bins, resulting in 2048 bins covering the 0-5 Hz frequency range. Next, we distinguished the very low frequency (0.01-0.08 Hz) and individual respiratory, and cardiac FFT power frequency ranges from the global and voxel-wise MREG periodograms. We achieved this by applying fslroi after specifying the relevant individual frequencies, and then calculated the summed power over each frequency range using the 3dTstat function as described in (20). The cardiorespiratory pulsation frequencies were also verified using the MR scanner respiratory belt (or anesthesia ETCO_2_ monitor Datex Ohmeda Aesthiva 5) and right index fingertip SpO_2_. For summary results, see Table 1. Finally, we performed a paired statistical analysis of the mean power maps between awake and sleep states using FSL-TFCE randomise (threshold p < 0.05). We applied brain structural segmentation using FSLeyes edit mode, and used fslcc to compute the correlation coefficients between power and speed analysis results.

### Flow speed verification process

We confirmed flow speed using a phantom consisting of a fresh pineapple (having a striated tissue structure), which had been prepared by drilling a 7 mm diameter hole through the core, through which we pumped water with a peristaltic pump (Watson Marlow 313S) via an elastic tube arising from outside the Faraday cage. The phantom imaging began with a 1 min baseline recording, followed by recordings at three flow speed stages (10, 20, and 30% of max speed, i.e., 400 rpm), each lasting 1 min. For an illustration of the phantom measurement and the ultrasound speed verification protocols, c.f. Supplementary Figure 3.

After scanning the pineapple, we preprocessed the MREG data by subtracting the mean of baseline signal from all other flow-rate stages. Then, we aligned the MREG data with anatomical T1 & T2 images after motion control and other pre-processing steps, whereupon we performed sparse and dense optical flow analyses matching those performed on subject data. These data were analyzed with a resolution pyramid of scale 3, because the peristaltic pulsation frequencies were faster than cardiovascular pulses.

## Results

### Flow speed verification result

Every stage of the three flow speed stages (i.e. 10, 20, and 30% of the maximum pump speed) produced a peak in the power spectrum, as seen in Figure 2A and Supplementary Figure 4. For each successive pump acceleration, the power and frequency of the MREG pulsation spectra also increased, accurately reflecting the pulsation power increase inside the pineapple phantom. However, from the power spectra, we find a clear aliasing in the fastest stage, where the pump frequency exceeded the Nyqvist theorem cutoff in our 10 Hz sampling rate. Therefore, we did not further analyze the results from the third speed stage.

**Figure 2.**
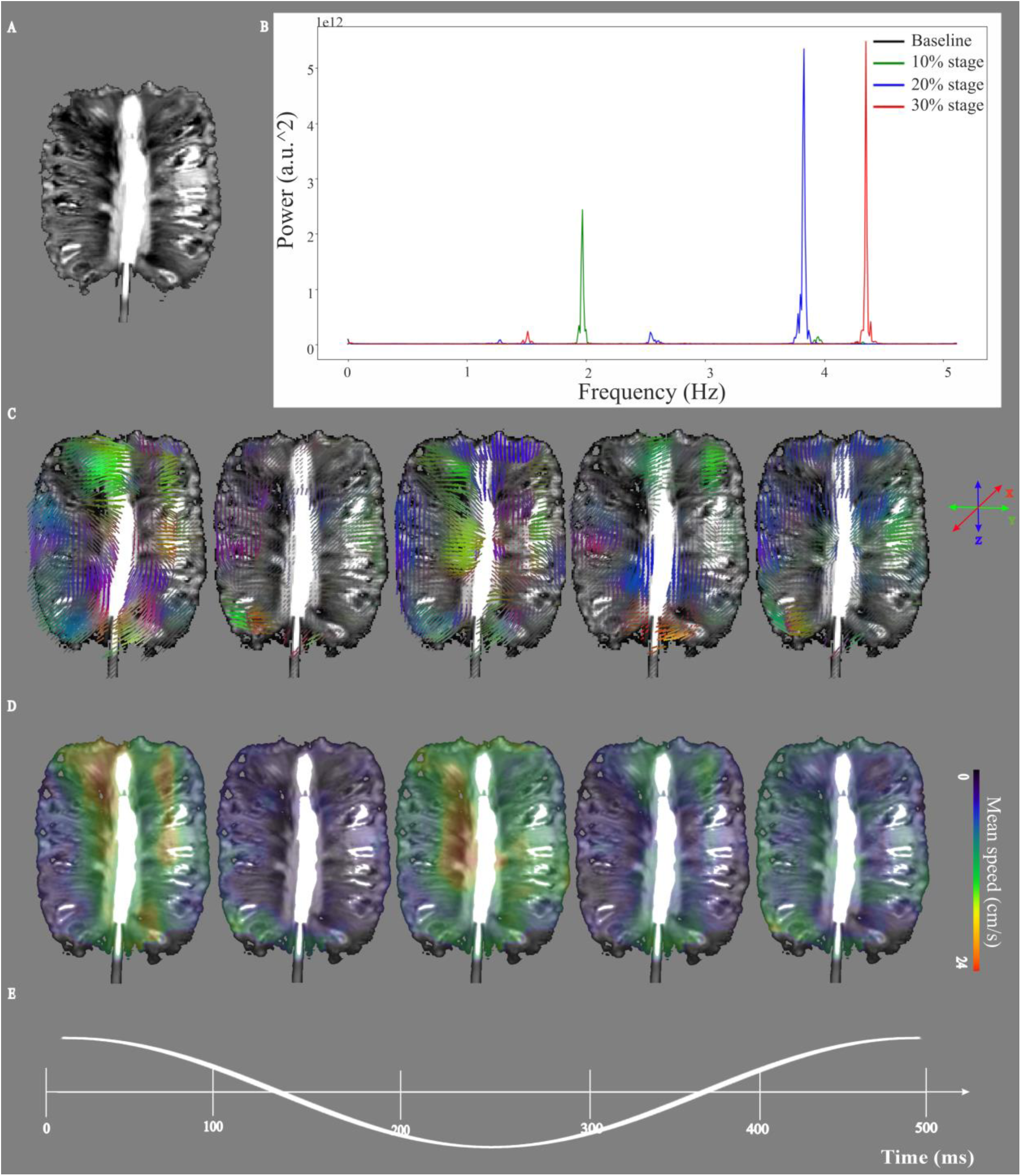
Mean velocity of optical flow analysis on stage 1 flow speed through the pineapple. (A) Imaging of water flow through a pineapple phantom using the same MREG settings as for the study subjects. (B) An FFT power spectrum of the MREG data accurately depicts the water pump frequencies that were then used for sub-sequent band-pass image filtering at the four stage of flow speed inside the pineapple. (C) The 3D directional V maps of stage 1 water flow impulse propagation analyzed in five-time segments over the averaged 0.5 s cycle. (D) Speed **v_s_** analysis of stage 1, over the five-time segments of the cycle. (E) Representative time domain signals of stage 1.

We also found a direct relation between the power and speed inside the porous medium of the pineapple. The results of a 3D V analysis of stage 10% of the maximum pump speed can be seen in Figure 2C; for results of the 20% maximum pump speed, see Supplementary Figure 5. For results for the sparse optical flow analysis of the same pump speeds, see Supplementary Figures 6-7. We emphasize that the speed vectors in the directional measures align to the structural streaks inside the pineapple flesh (Figure 2C). The flow magnitude maxima naturally align along the borehole direction. Interestingly the speed shows a double-peaked maxima during the peak and nadirs of the impulse cycle, which closely matches typical findings inside the human brain (Figure 2D).

Table 2 presents a comparison of results from the ultrasound measure (as a ground truth), along with sparse and dense optical flow. The results from the Doppler ultrasound analysis are detected by the maximum flow speed. To obtain a robust measure of optical flow for the 1-minute data époques, we calculated the mean of all the signal maxima across a cross-sectional area of the phantom, located in front of the flow input.

**Table 2.** Comparison between different analysis methods for measuring optical flow at different maximum speeds detected by Doppler ultrasound just outside the point of entry into the pineapple. The dense method, which detects all movement changes, is more accurate in detecting the maximal speeds than is the sparce method, which searches only a signal peak movement.

| Measurement | 10% stage (cm/s) | 20% stage (cm/s) |
| --- | --- | --- |
| Ultrasound | 22 | 33 |
| Dense optical flow | 23.64 | 34.82 |
| Sparse optical flow | 13.15 | 37.81 |

### Sleep staging results

The AASM-based sleep classification performed by clinical neurophysiologists yielded the following percentage scan times as a function of arousal state during the sleep scans: Awake 10 %, N1 stage 43 %, N2 stage 43 %, N3 stage 1%, artifacts 2 %. Therefore, in total, 87 % of the selected sleep scan époques were scored as NREM sleep. Corresponding values for awake scans were: Awake 99 %, N1 stage 0.5 %, and artifacts 0.5 %.

### Human Brain Cardiovascular impulse velocity

Overall, the cardiovascular impulse propagation speeds (**v_s_**_card_) in whole brain were the fastest of the three physiological brain pulsations, with an average speed of 3.27 ± 0.63 cm/s and maximum of 9.39 cm/s for the awake state: For the NREM sleep state, the average speed was 2.98 ± 0.67 cm/s and the maximum 7.41 cm/s. The spatial 3D velocity of cardiovascular pulsation (V_card_), along with changes in the direction of cardiac pulse propagation (<u>v</u>_card_) are presented in Figure 3A; for more comprehensive, 3D dynamic views of the speed changes, see Supplementary video S1.

**Figure 3.**
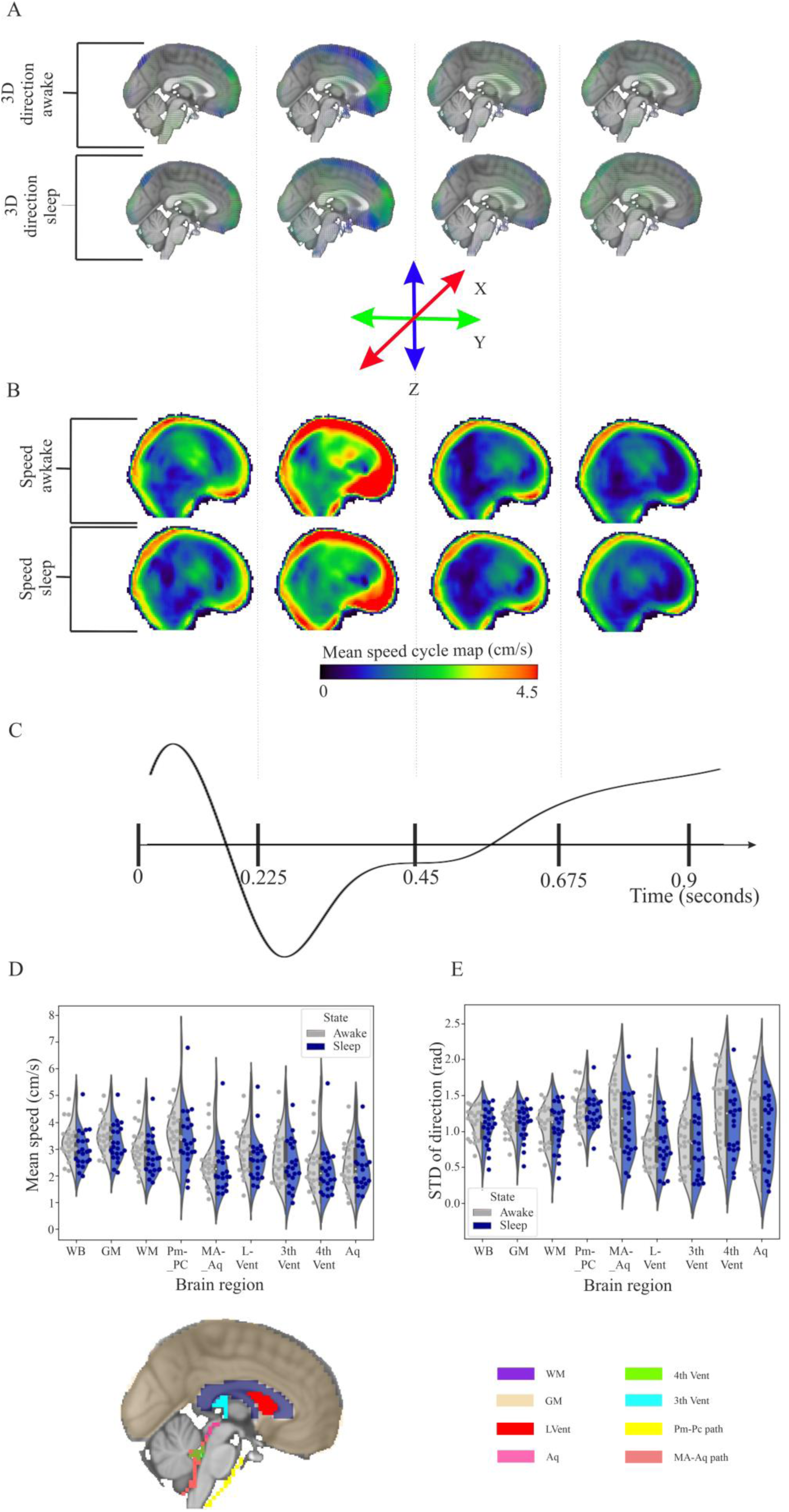
Optical flow analysis of mean cardiovascular brain pulsations in awake and sleeping subjects. (A) The 3D directional V_card_ maps of the cardiovascular impulse propagation were analyzed in four separate time segments over the averaged 0.9 s cardiac cycle. (B) Mean **v_s_**_card_ of awake and sleeping subjects, over the four-time segments of the cardiac cycle. (C) Representative time domain signal of the anterior cerebral artery. (D) The mean **v_s_**_card_ and (E) the STD of <u>v</u>_card_ in study subjects in both arousal states over the whole brain and in segmented brain structures, illustrated in the sagittal and axial planes of the MNI brain; Whole brain (WB), grey matter (GM) and white matter (WM), median aperture (MA), pontomedullary cistern (PM), pontine cistern (PC), all CSF ventricles (Lateral, III, IV) and aqueduct (Aq) are presented. For more details, see a 3-plane dynamic video of wave speed in Supplementary video S1.

The average 0.9 s cardiovascular cycle shows maximal impulse **v_s_**_card_ at the arrival of every heartbeat to the ACA, approximately 0.3 s after the cardiac R peak on ECG, as shown previously in control and Alzheimer’s disease patient populations (Rajna et al., 2021)(16). In the dense optical flow analysis, the arteries and cortical CSF spaces show the highest speeds. We show, however, that the mean **v_s_**_card_ (cm/s), and the inter-individual standard deviation (STD) of <u>v</u>_card_, in radians, varies between different brain regions of interest, as illustrated in Figure 3D-E.

After statistically comparing V_card_, neither the direction vectors <u>v</u>_card_ nor the magnitude **v_s_**_card_ significantly differed in NREM sleep versus awake state. Overall, the mean **v_s_**_card_ speeds were somewhat higher than 3 cm/s in grey and white matter and in the pre-pontine CSF around the basilar artery, as compared to the 2 cm/s for CSF spaces in the ventricles and around the cerebral aqueduct (Figure 3D). The mean STD of the directionality for all compartments was close to 1.2 radian, albeit somewhat lower in CSF spaces, but with higher variance between subjects. Furthermore, neither the mean speeds nor the directional variance of the cardiac pulse in the ROIs differed across arousal state (Figure 3E).

### Respiratory brain pulsation velocity

The mean respiratory speed **v_s_**_resp,_ in the awake brain was 0.26 ± 0.07 cm/s, with a maximum of 1.1 cm/s, versus a mean **v_s_**_resp_ of 0.31 ± 0.01 cm/s and maximum of 1.11 cm/s in the sleep state. The pattern of 3D respiratory pulse propagation in the brain was spatially more complex than for cardiac propagation, c.f. Figure 3A-4A, which is best appreciated from viewing the 3D dynamic Supplementary videos S1-3. The respiratory velocity vectors (V_resp_) illustrate this complexity of pulsation directions in brain tissue. Here, the magnitude of the velocity (**v_s_**_resp_) is as high as 0.8 cm/s in the basal CSF areas and in brain structures proximal to the basal CSF volumes and conduits, and likewise in cerebellum, brain stem, and, interestingly, in the hippocampus, a key brain structure for memory consolidation. Similarly to our previous findings in healthy controls (17), the **v_s_**_resp_ increased in a similar bi-phasic manner over the respiratory cycle as seen in the phantom experiment. The **v_s_**_resp_ reached its maximum during the initial quarter of exhalation and the initial quarter of inhalation, as determined by the mean MREG signal from the IV ventricle ROI (see also Supplementary video S3).

**Figure 4.**
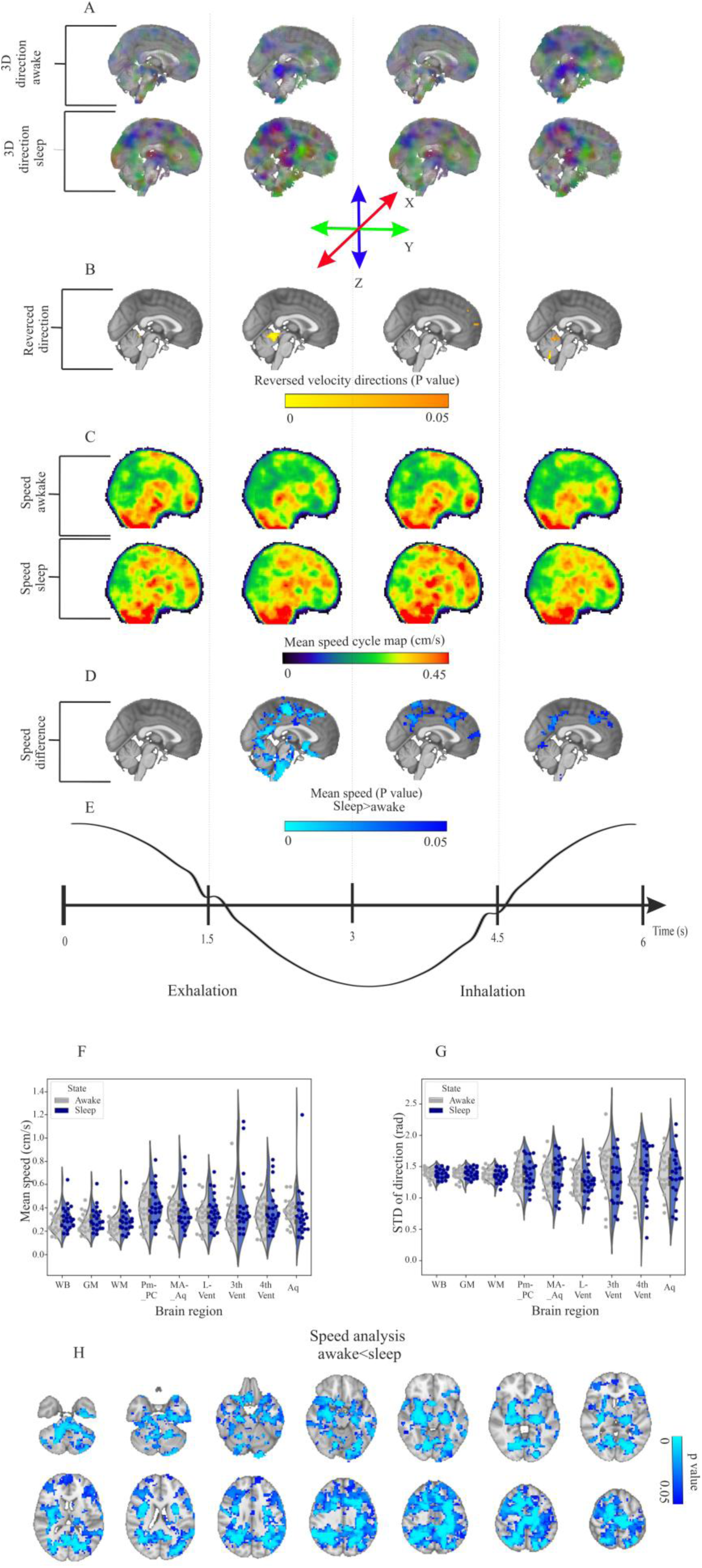
Mean respiratory speed increase during sleep throughout the neocortex and subcortical regions. (A) Mean directional V_resp_ for awake (n=22) and sleep (n=22) subjects. (B) Reversed <u>v</u>_resp_ in paired statistical analysis between awake and sleep groups (randomise FWER correction, p<0.05). (C) Mean **v**_sresp_ over a complete respiratory cycle, which was resampled to 6 s lengths for the awake and sleep groups. (D) **v**_sresp_ differences (randomise FWER correction, p<0.05). (E) Representative time domain signal of the IV ventricle. (F) Distribution of mean V_resp_ between groups over a complete respiratory cycle in different brain regions, and (G) STD of the direction. (H) Summed speed differences between awake (n=22) and sleep (n=22) groups (sleep>awake, FSL randomise, FWER correction, p<0.05) over the complete respiratory cycle summarized in one frame. See also Supplementary video S3, showing dynamics of the speed changes across the respiratory pulse cycle, with evident differences between arousal states in 3D.

In comparison to the awake state, the **v_s_**_resp_ increased significantly (FWER correction, p<0.05) in the NREM sleep condition. This increase was most significant during the latter half of exhalation and in early inhalation in comparison to awake state, while there was an increase during the first half of the exhalation encompassing a small area, as seen in Figure 4D. The **v_s_**_resp_ increased in the NREM state close to the sagittal sinus, in frontoparietal cortical areas, and especially in subcortical white matter, upper bilateral putamina, thalamus, and hippocampal areas, as well as in the lower brain stem.

The inspirational speed increases occurring in NREM sleep had a more circumscribed spatial distribution and lower magnitude compared to the exhalation changes (Figure 4D). During middle inspiration, the respiratory impulse propagation speed again increased in subcortical white matter in the upper frontal and parietal cortical brain areas, while the basal flow speeds fell back to awake levels (Figures 4D/H). There were directional <u>v</u>_resp_ changes in the propagation of the respiratory pulse in the cerebellum, the CSF space below the tentorium, and in small frontotemporal cortical areas (Figure 4B).

Figure 4F depicts the mean structural **v_s_**_resp_, which tended to be lower in the brain ROIs compared to the CSF areas. Interestingly, the directional variation in the tissue fell very tightly within about 1.4 radians, with very little population variance. On the other hand, the CSF spaces showed much more directional variation, especially in the III, and IV ventricles and in the aqueducts (Figure 4G).

On average, the speed of the respiratory pulse propagation increased significantly across the brain over the whole cycle, c.f. Figure 4H and Supplementary video S3. The mean **v_s_**_resp_ over the whole brain was about 28.5% higher over the whole brain in the sleep state compared to the awake state. Overall, the summed results over the whole cycle (Figure 4H) showed increases in the cerebellum, temporomesial hippocampal areas, in the thalamus, and in periventricular WM ROIs, posterior DMN, and sensory cortices.

### Vasomotor brain pulsation velocity

The propagating vasomotor waves arising from the PCC, which is a component of the default mode network (DMN_PCC_), displayed a distinctly sinusoidal MREG temporal signal pattern (Figure 5E). The waves were of low frequency (< 0.08 Hz) and low speed (mean **v_s_**_vaso_ was 0.058 ± 0.009 cm/s), and the measurable maximal speeds were around 0.204 cm/s, i.e. only 2.12% of the quantifiable maximal cardiovascular speeds. In the sleeping subjects, the mean speed was 0.06 ± 0.01 cm/s, and the maximum speed 0.253 cm/s. The vasomotor waves also presented a rich dynamic 3D spatial pattern of propagation, surpassing in complexity the cardiac pulsations, c.f. Figures 3A-5A. and Supplementary video S4. Like the respiratory speed magnitudes, **v_s_**_vaso_ also presented a bi-phasic speed pattern in BOLD signal up and signal down phases of the VLF signal. Also like the respiratory pulse, the **v_s_**_vaso_ maximized near the mean signal level at the early decrease and increase phases, with steepest MREG signal changes in DMN_PCC,_ (Figures 5A/C, Supplementary video S4).

**Figure 5.**
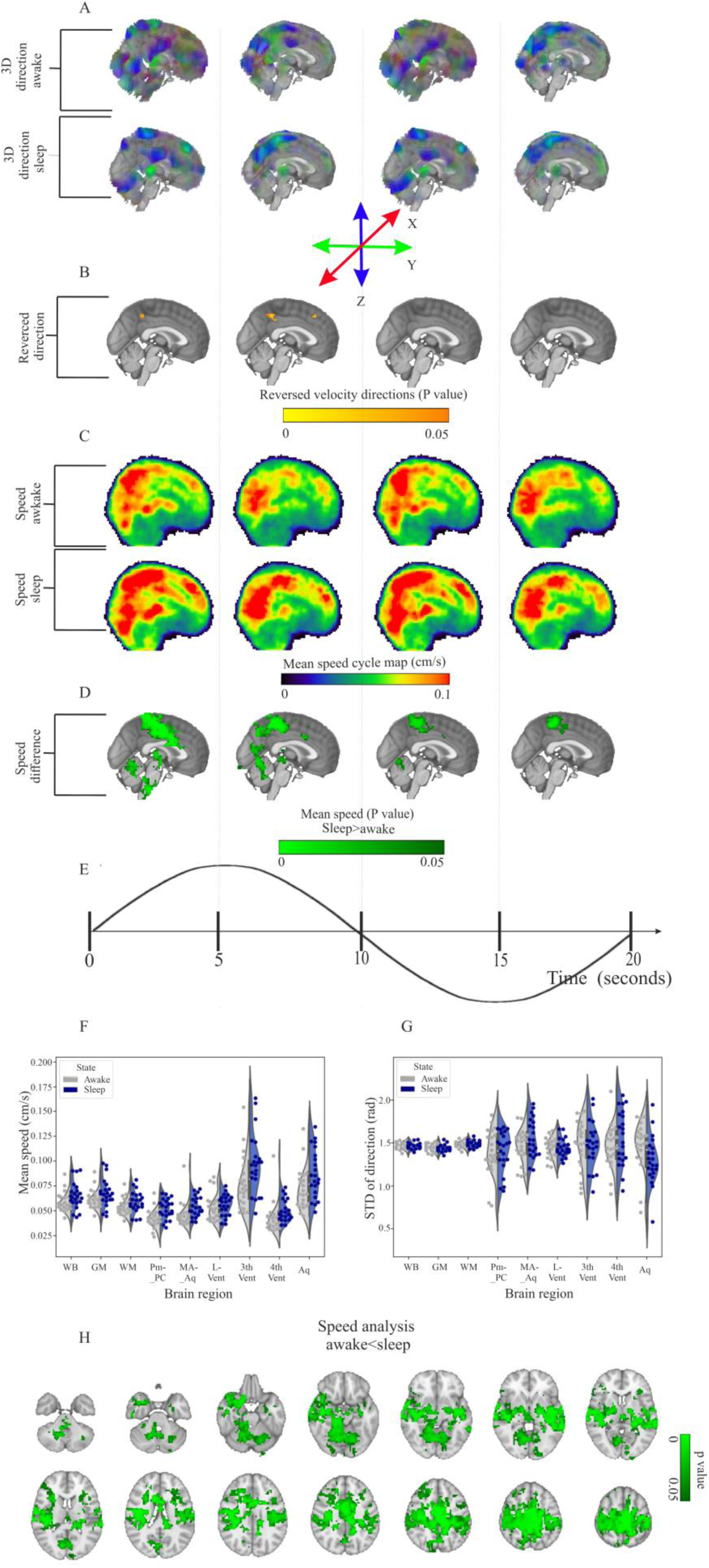
Mean speed of vasomotor brain pulsations increases during sleep. (A) Optical flow analysis of the mean physiologically awake (n=22) and sleeping (n=22) subjects, over a complete vasomotor cycle, and (B) reversed velocity direction <u>v</u>_vaso_ in a contrast of the awake and sleep groups (FSL randomise, FWER corrected, p<0.05). (C) Mean speed **v**_svaso_ of awake (top) and sleep (below) groups, over a complete vasomotor cycle, and (D) statistical difference in **v**_svaso_ speed between the groups at the four stages of the vasomotor cycle (sleep > awake, FWER corrected, p<0.05), see also Supplementary video S4. (E) Representative time domain signal of the PCC. (F) STD of <u>v</u>_vaso_ of awake and sleep subjects extending over a complete vasomotor cycle, and (G) STD of <u>v</u>_vaso_ between groups. (H) Fourteen axial slices showing temporal summation of significant increases in VLF vasomotor wave speed (sleep > awake, FWER correction, p<0.05). For dynamic speed changes in 3D, see also Supplementary video S4.

The sleep group also showed significant increases (FWER correction, p<0.05) in the of the VLF vasomotor wave propagation by some 21.2 % extending over the whole brain. Across the whole cycle length, the summed speed increases occurred mostly in areas encompassing the sensory (auditory, sensorimotor, and anterior visual) brain cortices, bilateral thalamus, and the frontal cortices. Hippocampi and especially the cerebellum lying close to the IV ventricle showed right dominant speed increases in the sleep condition.

In ROI analyses, the **v_s_**_vaso_ were higher during sleep in the III ventricle and cerebral aqueduct, albeit with high-speed variation compared to other segmented brain structures (Figure 5F). The CSF spaces showed increased variance in the speed vector directionality, similarly to respiratory pulsations in comparison to the tight 1.4 radian directionality measured in brain structures (Figure 5G).

Dynamically, the higher **v_s_**_vaso_ in the sleep condition was most distinct during the 1^st^ quarter of the vasomotor wave cycle, extending over brain regions assigned to the task-positive network, mostly involving somatosensory and auditory cortices, and anteriorly in the V2 visual cortex. The **v_s_**_vaso_ increase during sleep extended from sensorimotor areas over the SMA into the midline cingulate gyrus, bilaterally in the pallidum, amygdala, and hippocampus, midline thalamus, and in right temporal areas. The right cerebellum, brain stem and IV ventricle also showed notably increased vasomotor wave propagation speed.

Towards the end of the averaged 20 s vasomotor cycle, a transient increase of the vsvaso tended to normalize in frontal and basal areas, but there remained an increase over the somatosensory areas and upper cerebellum during some three quarters of the complete VLF wave cycle (Figure 5H). There was a locus of continuous speed increase in the SMA/juxtapositional lobule in midline parasagittal areas; for dynamics, see the 3D video in Supplementary video S4.

### Power vs. speed of the physiological brain pulsations

We have previously detected significant power increases in all three physiological brain pulsations during sleep (20, 21). As the power of waves influences the flow speed inside porous media (27, 28) such as the brain, we further compared the spatial co-localization of the detected speed and power changes shown in Figure 6. First, the cardiovascular power does not markedly increase in NREM sleep and accordingly there is no speed increase in these fastest pulsations. However, we found a wide-spread spatial overlap between the results of speed and power analysis of the slower physiological pulsations in sleep. The spatial correlation coefficients between power and speed difference maps in respiratory and vasomotor sleep changes were r = 0.71 and r = 0.79, respectively. This indicates widespread covariance in overlapping regions. However, between the two arousal states, the power change had a wider spatial distribution compared to the speed changes. Furthermore, the spatial distribution of the speed changes also tended to increase inversely with the frequency of the pulsation, being strongest in vasomotor and virtually absent in cardiovascular pulsations.

**Figure 6.**
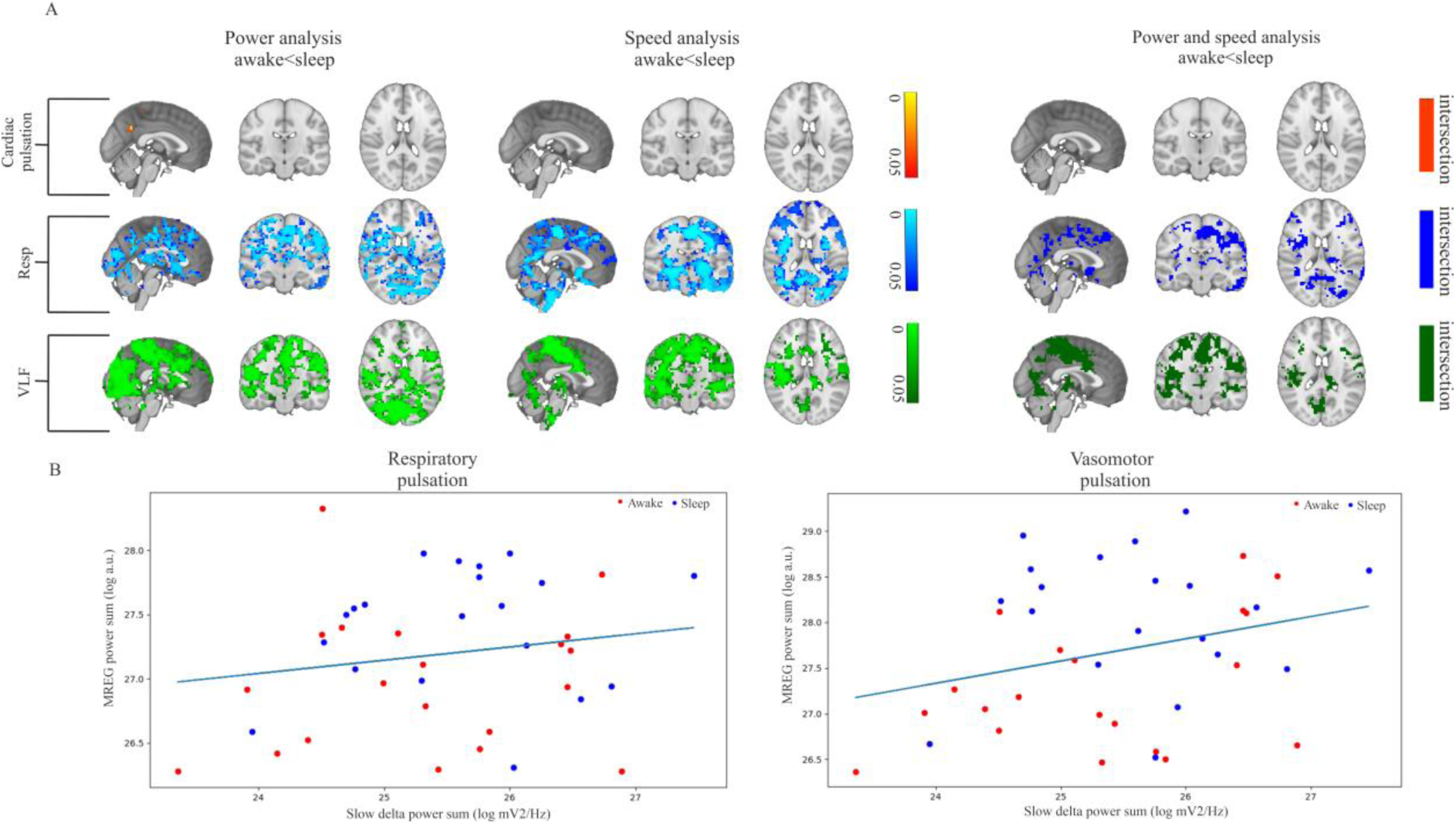
Power, speed and slow-delta power analysis of subjects in the two arousal states. **(A)** There were a few alterations evident in the power analysis of the cardiac pulsations in the precuneate cortex, however, we did not detect any alterations in the speed analysis for cardiac pulsation. On the other hand, we detected wide-spread and spatially overlapping increases in the respiratory and vasomotor pulsations in the sleep condition. **(B)** MREG power vs. slow Delta EEG power (EEG_S.Delta_) in each subject for respiratory and likewise vasomotor pulsations. The speed did not correlate significantly with the slow Delta power, as seen in supplementary Figure 8.

Neurophysiological delta power increases in mouse brain areas that are flushed with CSF, which indicates increased glymphatic solute transport during sleep (1). In human brain, physiological pulsation power likewise increases in brain regions showing an increase in the neurophysiological slow delta power during sleep (20, 21). A more recent study on a different population verified this finding and showed that the cardiorespiratory power and slow-delta powers are linearly correlating and differential in awake and sleep (35). In the present study, we further investigated whether the concomitant power increases in the physiological pulsations would likewise reflect slow delta EEG power changes. We found that the simultaneously measured slow delta EEG power increases that occurred in sleep correlated with the whole-brain power of vasomotor and respiratory pulsations, c.f. Figure 6B. The pulsation speed results did not have a clear association with the slow delta power, neither in whole brain, nor in the regions showing increased speed.

## Discussion

We first undertook a phantom experiment to confirm that MREG data measures water molecule movement in a porous medium. The phantom study furthermore confirmed that the dense optical flow analysis is of superior accuracy of detecting water molecule movement within a specified frequency band from the ultrafast 3D MREG data. Applying the dense optical flow analysis, we then proceeded to analyze in dynamic MREG data the sleep-related changes in human brain CSF propagation speeds, non-invasively and in the absence of aliasing. We detected significant increases in excess of 20% in the regional speed of both vasomotor and respiratory brain pulsations in NREM sleep vs. awake state; these increases were evident in specific phases of the pulsations, while the sleep-induced changes in cardiovascular impulse speed were unaffected by sleep. Importantly, in contrast to previous studies on the effects of pathology on pulsations speeds, there were only minimal changes in the direction of the velocity of the pulsations induced by natural NREM sleep. The areas of overall increase in speed were also spatially overlapping with the power increases both in vasomotor and respiratory pulsations.

### NREM sleep promotes pulsation speed as measured with optical flow

Tracer studies in animals have shown that brain solute convection and clearance increases during sleep (1, 47). Especially during NREM sleep, slow arterial vasomotor waves increase in speed, which in turn enhances fluid flow and drives perivascular CSF space volume changes (48). Physiological slowing of the brainstem *Locus coeruleus* noradrenergic neurons drives the sleep related arousal changes with potent VLF 0.02 Hz oscillations in norepinephrine levels (1, 47, 49). Human BOLD studies have also verified the occurrence of similar widespread vasomotor changes driven from the vicinity of the cholinergic nucleus basalis of Meynert (50), with additional contributions from sympathetic drive, especially in NREM light sleep (51, 52). Fultz et al. first described the connection between CSF flow fluctuation, BOLD signal changes, and EEG delta power across the sleep-wake cycle (53). That seminal work showed that the CSF flow BOLD signal in the IV ventricle followed increasing EEG delta wave activity, while being anti-correlated with the cortical gray-matter BOLD signal.

Previous work has also related EEG delta power during sleep to increased brain solute transport in mice (47), lending further support to the critical importance of brain fluid clearance also to neuronal activity. Picchioni et al. suggested that, in addition to the triggers arising in the CNS, there seems to be an autonomic pathway for regulating CSF flow, in a manner highly dependent on respiration rate (52). As such, we see a need for better elucidation of the causal relationships between physiological pulsations and arousal-dependent electrophysiological signals. In a phase contrasted MRI study, Dreha-Kulaczewski et al. showed that inspiration was the main promoter of CSF flow, with only small effects mediated by cardiac drive (10). Upon voluntary deeper and forced breathing, there was an increase synchronization of venous flow dynamics and CSF flow in human brain (54), and in another study, deep breathing increased CSF flow velocity (55, 56).

It is well-established that upper airway resistance, along with pulmonary and intrathoracic ventilation increase during sleep (57–59); these changes may have the net effect of promoting the pumping of CSF through the perivenous spaces in brain. This flow-enabling function may manifest as increased respiratory pulsatility in MREG. Indeed, our previous MREG results also indicated that the respiratory brain pulsation power increases as a function of sleep depth (20, 21). Another BOLD study showed that voluntary deep inspirations also increased the magnitude of CSF pulsations, and evoked widespread BOLD signal changes in cerebral cortex (52). In the present study, the sleep-related increases in speed and power of both (respiratory and vasomotor) propagating physiological brain pulsations occurred in same sensory cortical areas. This result supports the proposition that not only VLF vasomotor waves, but also respiratory brain pulsations may be drivers for the increased CSF solute transport occurring in sleep.

### On the physiology of the brain pulsation speed changes

Normal, unforced inhalation induces an inflow of CSF into the intracranial space (54), during which the MREG signal also increases in the IV ventricle, with converse effects during exhalation, c.f. Figure 4, Supplementary video S3. The maximal flow speeds occur at the midpoints of exhalation and inhalation, but sleep-induced increases in the respiratory **v_s_**_resp_ speed main occur when the respiration cycle switches from exhalation to inhalation. After the peak, the speed gradually declines until the end of inhalation. At this turning point of the respiratory cycle, the transit speeds of red blood cell (RBC) decline by 25-33 % in brain surface arterioles and venules of mice (8). This declining perfusion could be due to a transient peak in CSF pressure, slowing fluid in/outflow, coincident with the nadir in **v_s_**_resp_ seen in the present study. As such, the reduced **v_s_**_resp_ during sleep may stem from CSF inflow, and/or venous blood outflow. Further work on mechanistic and microscopic level are needed to disentangle these two flow effects.

Like respiration, the speed of vasomotor pulsations also maximizes in the cerebral cortex at the moment of steepest MREG signal change observed in the tissue DMN_PCC_ trigger point (Figure 5, Supplementary video S4). The increased vasomotor VLF propagation speed during sleep maximizes in both spatial spread and speed at these maximal speed moments, (Supplementary video S4). This indicates that the maximal propagation speed of the vasomotor waves occurs at the beginning of vasoconstriction of precapillary arterioles/arteries, but also occurs strongly during the early vasodilation phase, with both effects driving the vasomotor waves. Given the recent discovery of perivascular volume pulsations during NREM sleep (48), these vasomotor speed increases suggest involvement of altered perivascular flow and volume propagating in the brain tissue as waves.

Even though cardiovascular pulses primarily drive the periarterial space CSF flow in the murine sub-arachnoid space and cortex (4), we did not find any differences in cardiovascular pulsations in the present contrast of awake versus sleep conditions in humans. There were no sleep-related speed increases of the relatively mono-phasic cardiovascular impulses, nor was there any sign of a directional change. This further suggests that vasomotor and respiratory pulsations together promote solute transport in the human brain during NREM sleep. However, taken together all the three pulsation results, the finding that power increases are needed for pulsation speed to increase, implies that porous media fluid physics apply in the human brain hydrodynamics.

### Pathology vs. physiological sleep induced changes in pulsations speed

Patients with Alzheimer’s disease showed changes in cardiovascular pulsation speed, with some regions exhibiting reversed flow direction compared to healthy controls (16). Importantly, physiological processes like sleep do not inherently change speed nor alter the directionality of any of the cardiovascular pulsations. This suggests that the reversed pulse propagation in Alzheimer’s disease may be more reflective of inverted flow impulse patterns rather than normal physiology.

In patients with epilepsy, current methods have revealed an abnormally low speed of respiratory pulsations compared to healthy controls, particularly during the early phase of exhalation, which broadly affects the brain (17). Additionally, signal variation and the power of respiratory pulsations are both elevated in these patients relative to healthy controls (25). Interestingly, during normal sleep, both pulsation speed and power increase, whereas these changes are reversed in epilepsy.

In Alzheimer’s disease and epilepsy, the brain may experience some degree of homeostatic compensation for the pathological changes. It is well-established that sleep deprivation impairs glymphatic function and contributes to the accumulation of amyloid beta in the brain (60, 61). This highlights the potential importance of adequate sleep-in enhancing brain metabolite clearance, which been shown to reduce the risk of developing Alzheimer’s disease (62–64) and to alleviate symptoms of epilepsy (65).

### Correlation of speed with pulsation power and slow delta EEG power

We found spatially overlapping patterns of altered **v_s_**_resp_ and **v_s_**_vaso_ pulsation speeds during NREM sleep compared to waking. These overlapping regions exhibited increased power for both pulsations, which was associated with elevated EEG delta power (see Figure. 6). Moreover, we replicated the linear correlation of the power and found no significant correlation in speed with respect to slow delta EEG-power, as seen in supplementary Figure 8 (35). Regarding causality, the slow delta wave power may facilitate the propagation of respiratory waves, leading to faster movement. Alternatively, increased delta power could be a consequence of enhanced solute transport, potentially influencing EEG power by expanding the interstitial space, which could affect neuronal transmission. The elevated delta power may result from increased exchange between interstitial fluid (ISF) and cerebrospinal fluid (CSF) in the perivascular spaces during sleep (1), with additional contributions from noradrenergic mechanisms as described previously (49).

Human sleep experiments have shown that slowly emerging K-complexes in N2 sleep, and likewise during controlled deep breathing, cause similar CSF inflow changes, which couple to widespread brain BOLD signal changes (52). In general, a higher amplitude wave will have more energy, and can thus can travel faster through a porous medium, depending on the viscous properties of the tissue (26–28). The stronger vasomotor waves that occur in NREM sleep (20, 48, 50, 53) could relate to an increased flushing between ISF/CSF in the perivascular spaces. We suppose that this phenomenon might alter the local electrolyte concentrations, and drop overall EEG activity towards delta power in relation to increased water entry into the interstitium.

### Technical aspects, strengths, and limitations of MREG mapping

The phantom study cleared showed that water inflow to a porous medium is precisely reflected into the MREG signal data, c.f. Figure 2 and Supplementary Figure 5. Increased flow speeds and pump frequency are unequivocally reflected by MREG mapping of the phantom. Moreover, to dense optical flow, the increased pulsation power and frequency correlated with increased flow speeds inside the porous pineapple tissue. This identical phenomenon was also detected in the perivascular CSF tracer studies in living animals in a recent paper by Nedergaard group, where the increase in vasomotor wave frequency promoted faster glymphatic CSF flow and clearance (66).

We find that the same biphasic pulsation flow patterns detected in the phantom pulse cycles are also present in the living human brain. The lacking power increase in cardiac frequency in sleeping human brain is related to the lacking speed changes, which is also evident in the stable results within the 1 min periods of constant pumping rates in the phantom experiment. In the human brain, dense optical flow analysis of MREG data indicated maximal cardiovascular pulsation speeds of 9.39 cm/s, which is close to the range of CSF flow speeds (5-8 cm/s) that are driven by the cardiovascular pulses in normal awake breathing (67–69). The present dense optical flow MREG results of 3.27 ± 0.63 cm/s and maximum flow of 9.39 cm/s for the awake state are in good agreement with the earlier reports.

In the confines of the aqueduct, CSF flow speed ranges from 2.8 mm/s to a maximum of 11 mm/s (70). Also, the tissue arteriolar erythrocyte speeds range up to 22.2 mm/s, but increase from < 1 mm/s at the capillary level to as much as 5 mm/s in venules (8). The present tissue speeds of **v**_svaso_ are in line with the slow venule speeds, insofar as **v**_sresp_ largely reflects flow in larger veins and perivenous spaces (8).

In the future, still faster MREG methods, preferably with higher spatial resolution, may emerge for better comparison with invasive contrast studies. Velocity encoded and diffusion-weighted MREG promises to reveal an improved depiction of the status of intracerebral hydrodynamics. Our present comparison of the established sparse optical flow method with the dense optical flow method showed similar performance for detecting cardiovascular pulsations. However, the dense optical flow method outperforms the sparse method for slower pulsations, as clearly shown in Supplementary videos S1-2, Figure 3, and Supplementary Figure 1.

In the brain parenchyma, the respiratory pulses predominate around the outflowing veins, and cardiovascular impulses are strongest around the inflowing arteries, where they augment respiratory effects (4, 8). The venous BOLD effects driven by susceptibility effects dominate both the respiratory and vasomotor frequencies, and spin-phase changes reflect the faster cardiovascular impulse changes in the MREG signal. In the CSF spaces T2 phase effects are the main signal source (71, 72): for detailed information on the nature of the MREG signal, (44). The ultrafast 10 Hz 3D whole brain sampling in MREG has an inherent trade-off with spatial resolution; however, the present 3 mm cubic voxels still offer considerable, spatially precise anatomic detail. Nonetheless, compared to single slice pcMRI, such 3 mm voxels impart significant partial volume effects, with signal impingement from neighboring blood vessels, CSF space, and brain tissue. Thus, MREG gives weighted average results for speed over the multiple vessels within each voxel, and are thus less sensitive to high maximal flow speeds within a single artery/arteriole, especially in the CSF/tissue interfaces with their opposing flow directions.

Intrathecal tracer studies have shown that cortical glymphatic mechanisms for solute transport in the 400 mg, lisencephalic mouse brain remarkably resemble those in the 3000x larger, convoluted human brain (73–75). Non-invasive methods for studying the driving forces of whole brain fluid clearance in both experimental animals and humans, are emerging, but not yet firmly established. Given the present lack of methods for following the direct effects of individual physiological pulsations on tracer kinetics, we opted to rely upon indices of how the physiological pulsations affect ultrafast T2* weighted water proton signal dynamics.

Falling asleep in the scanner is not easy, as lying on the scanner and its noise can cause discomfort, affecting sleep quality. Indeed, not all scanned data was usable due to lacking sleep époques. In addition, some EEG recordings were unusable due to excessive motion, imperfect electrode contact, or sweating. We decided to use 5-minute époques of continuous data that contained the highest amount of EEG-confirmed sleep, which necessarily contained a mixture of N1 and N2 sleep states. For similar reasons, we could not study acute effects of sleep onset. Although deemed healthy by self-report, we had not specifically screened the subjects for sleep patterns, sleep history, or sleep disorders. Our study in a small population of healthy young subjects does not generalize for the whole population. Finally, to compile sufficient data, we combined datasets of morning and evening scans, which may have introduced confounds from circadian rhythm in addition to the rebound sleep effects.

## Conclusions

The pineapple phantom experiment confirmed that MREG data indeed directed water molecule movement in a porous media, and showed the expected relationship between increased pulsation power and frequency with increased average flow speed in the tissue. The main part of the study highlights distinct characteristics of the 3D flow velocities and patterns in human brain as driven separately by cardiovascular, respiratory, and vasomotor pulsations, and revealed complex differences between sleep and awake states. Cardiovascular pulsations maintained the highest speeds and stability, with minimal impacted from sleep. Respiratory and vasomotor pulsations, however, exhibited widespread and more complex spatial patterns, with increased propagation speeds during sleep relating to increased pulsation power, particularly in memory-related (hippocampal) and sensory-processing cortical regions. These findings indicate that sleep significantly enhances lower-frequency pulsations, which increases CSF flow speed in a manner expected to promote metabolic waste clearance in human brain.

## Supporting information

Supplementary Figure 8.jpg

## Acknowledgements

We would like to thank all study subjects for their participation in the study. We also thank Annastiina Kivipää and Matti Pasanen for subject scanning and data preprocessing, and Jani Häkli and others who otherwise participated in the project. We wish to acknowledge Jussi Kantola for data management and reconstruction of MREG data, the CSC – IT Center for Science Ltd., Finland for generous computational resources, and Prof. Paul Cumming of Bern University for comments on the manuscript.

## Data availability

Data is available upon request

## Code availability

For 3D optical flow analysis, please contact

For the statistical analysis, please contact.

## Supplementary material

Supplementary material is available online

## Abbreviations

V: propagation Velocity
v_s_: propagation velocity magnitude (speed)
<u>v</u>: propagation velocity direction in Cartesian coordinates
<u>v_x_</u>: speed in x direction
<u>v_y_</u>: speed in y direction
<u>v_z_</u>: speed in z direction.
EEG_S.Delta_: slow delta power extracted from EEG data.

