## Supplementary Materials for "Sleep increases propagation speed of physiological brain pulsations"

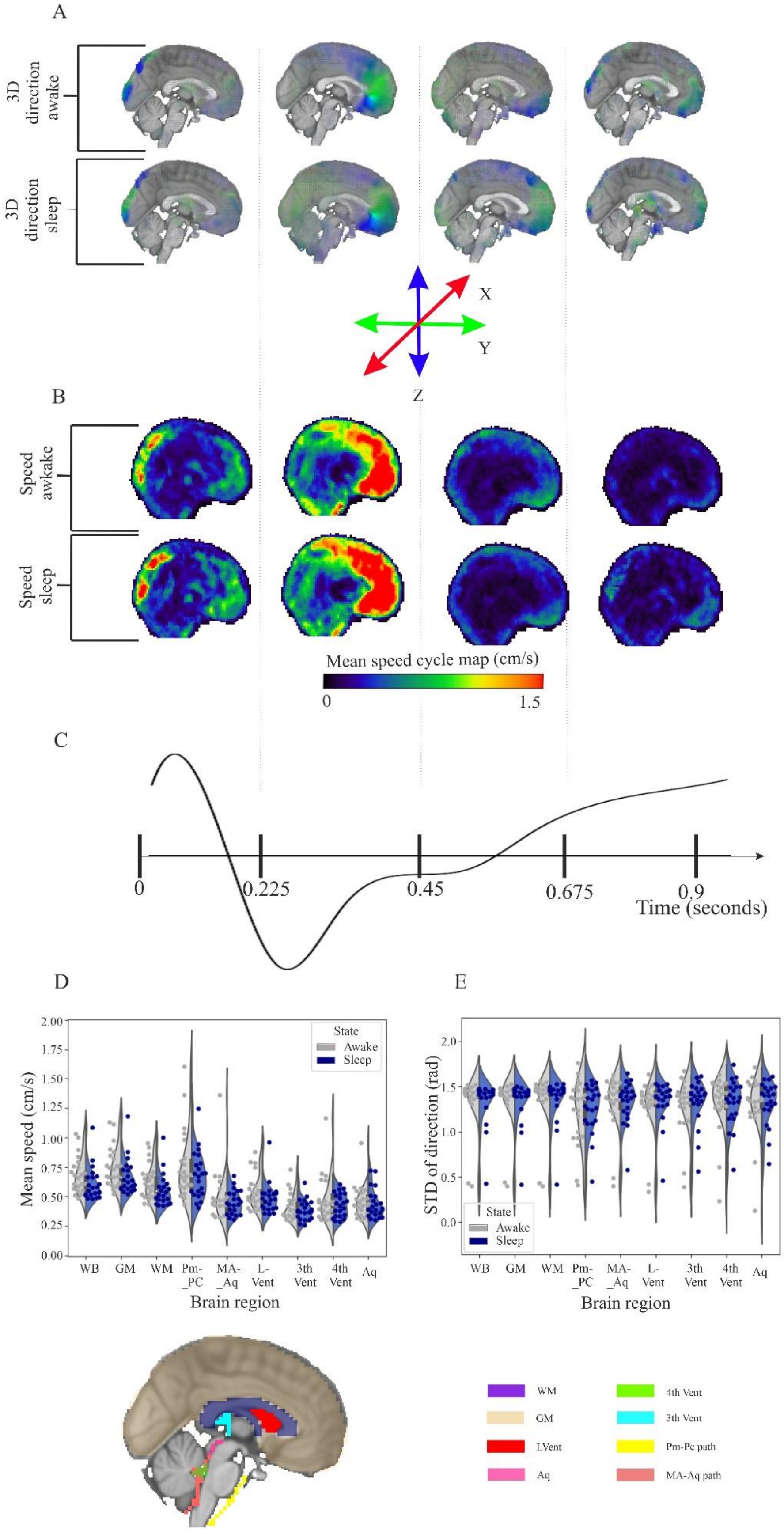

A
3D direction awake
3D direction sleep
X
Y
Z
B
Speed awkake
Speed sleep
Mean speed cycle map (cm/s)
0
1.5
C
0
0.225
0.45
0.675
0.9
Time (seconds)
D
State
Awake
Sleep
Mean speed (cm/s)
2.00
1.75
1.50
1.25
1.00
0.75
0.50
0.25
0.00
WB
GM
WM
Pm-_PC
MA-_Aq
L-Vent
3th Vent
4th Vent
Aq
Brain region
E
STD of direction (rad)
2.0
1.5
1.0
0.5
0.0
State
Awake
Sleep
WB
GM
WM
Pm-_PC
MA-_Aq
L-Vent
3th Vent
4th Vent
Aq
Brain region
WM
GM
LVent
Aq
4th Vent
3th Vent
Pm-Pc path
MA-Aq path

**Supplementary Figure 1. Optical flow analysis of mean cardiovascular brain pulsation in awake and sleep subjects.** (A) The 3D directional V maps of the cardiovascular impulse propagation is analyzed in four time segments over the averaged 0.9 s cardiac cycle. (B) Eigenvalue speed magnitude versus analysis of awake and sleep subjects, over the four-time segments of the cardiac cycle. (C) Representative time domain signal of the anterior cerebral artery. (D) The mean speed (E) and standard deviation (STD) of cardiovascular pulsations in study subjects in both arousal states over the whole brain, and in segmented brain structures illustrated in the MNI sagittal and axial planes; grey matter (GM) and white matter (WM), median aperture (MA), pontomedullary cistern (PM), pontine cistern (PC), all CSF ventricles (Lateral, III, IV) and aqueduct (Aq) are presented. For more detailed 3-plane dynamic video for biphasic speed wave, see Supplementary Video 1.

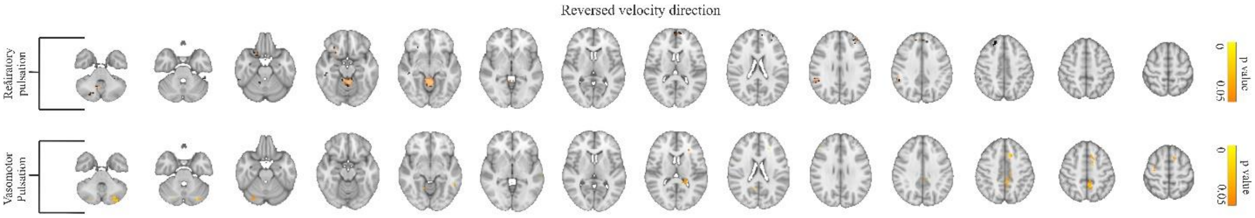


**Supplementary Figure 2**. **Significant velocity direction differences between awake and sleep states.** Reversed velocity direction in the comparison between awake (n=22) and sleep (n=22) groups (FSL randomise, family-wise-error-rate (FWER) correction, $p<0.05$)

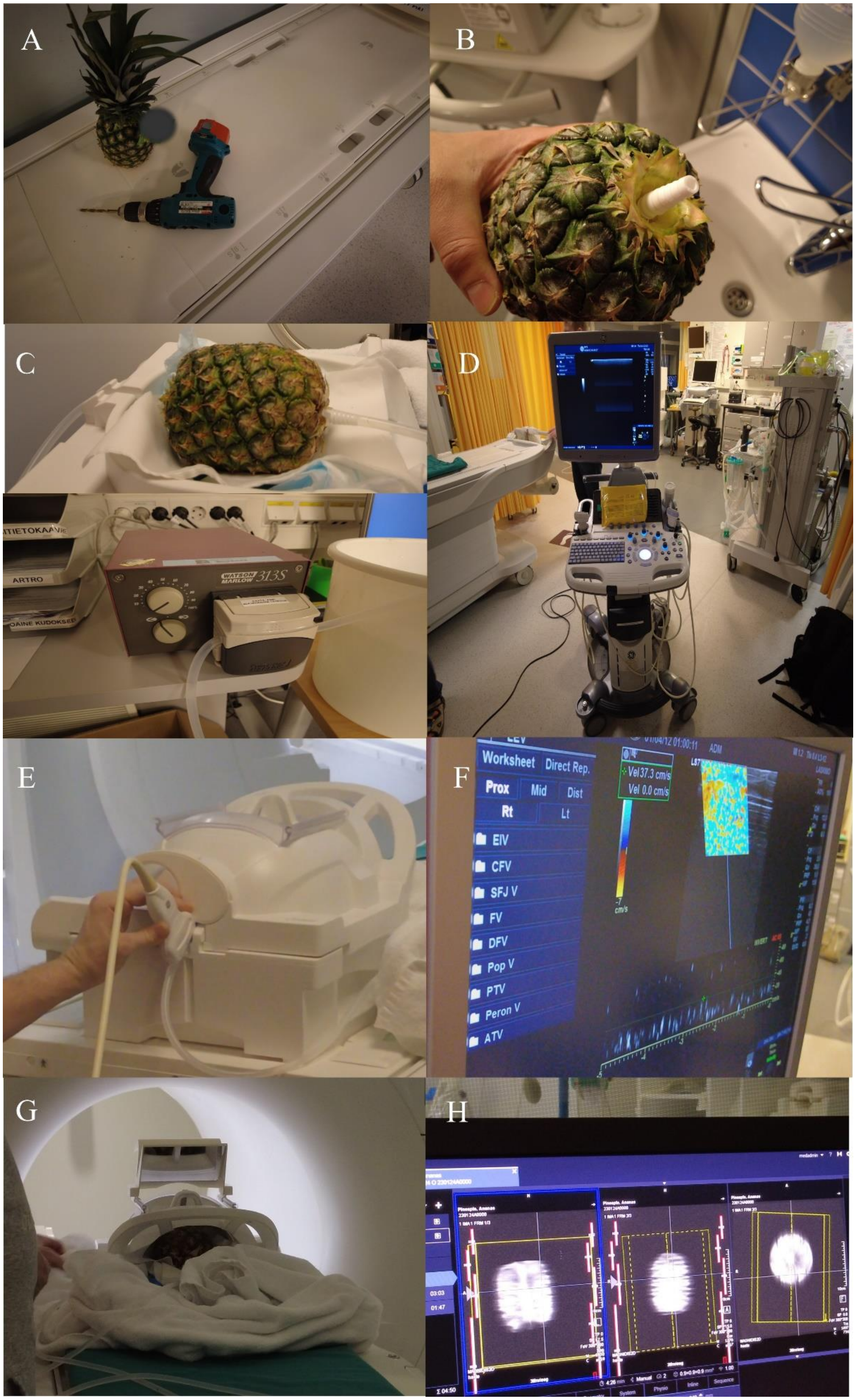
A
B
C
D
E
F
G
H
WATSON MARLOW 313S
ARTRO
Worksheet
Direct Rep.
Prox
Mid
Dist
Rt
Lt
Vel 37.3 cm/s
Vel 0.0 cm/s
EIV
CFV
SFJ V
FV
DFV
Pop V
PTV
Peron V
ATV

**Supplementary Figure 3. Scanning a pineapple protocol.** (A) We connected the peristaltic pump to the two plastic ends through elastic pipes. (B) Initialization of the ultrasonic meter. (C) Fixation of the meter peripherally to the point of flow input. (D) Initial measurement at (30% maximal pump speed). (E) Placement of the pineapple in the scanner. (F) Scanning the pineapple using the MREG sequence with the same settings as used for the study subjects.

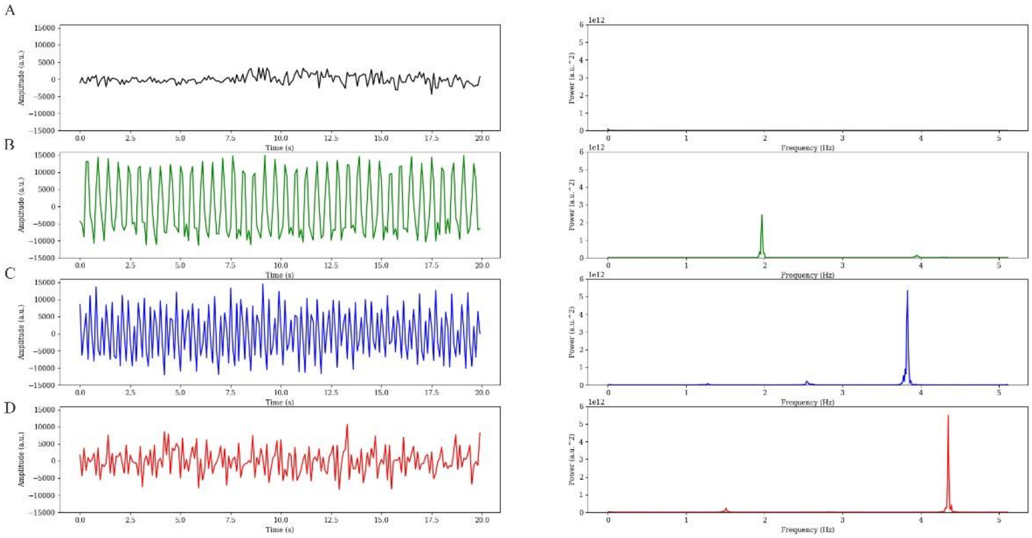


**Supplementary Figure 4. Time and spectral analysis of the four speed stages in the pineapple phantom study.** (A) Baseline signal. (B) 10% speed of the maximal pump speed. (C) 20% of the maximal pump speed. (D) 30% of the maximal pump speed.

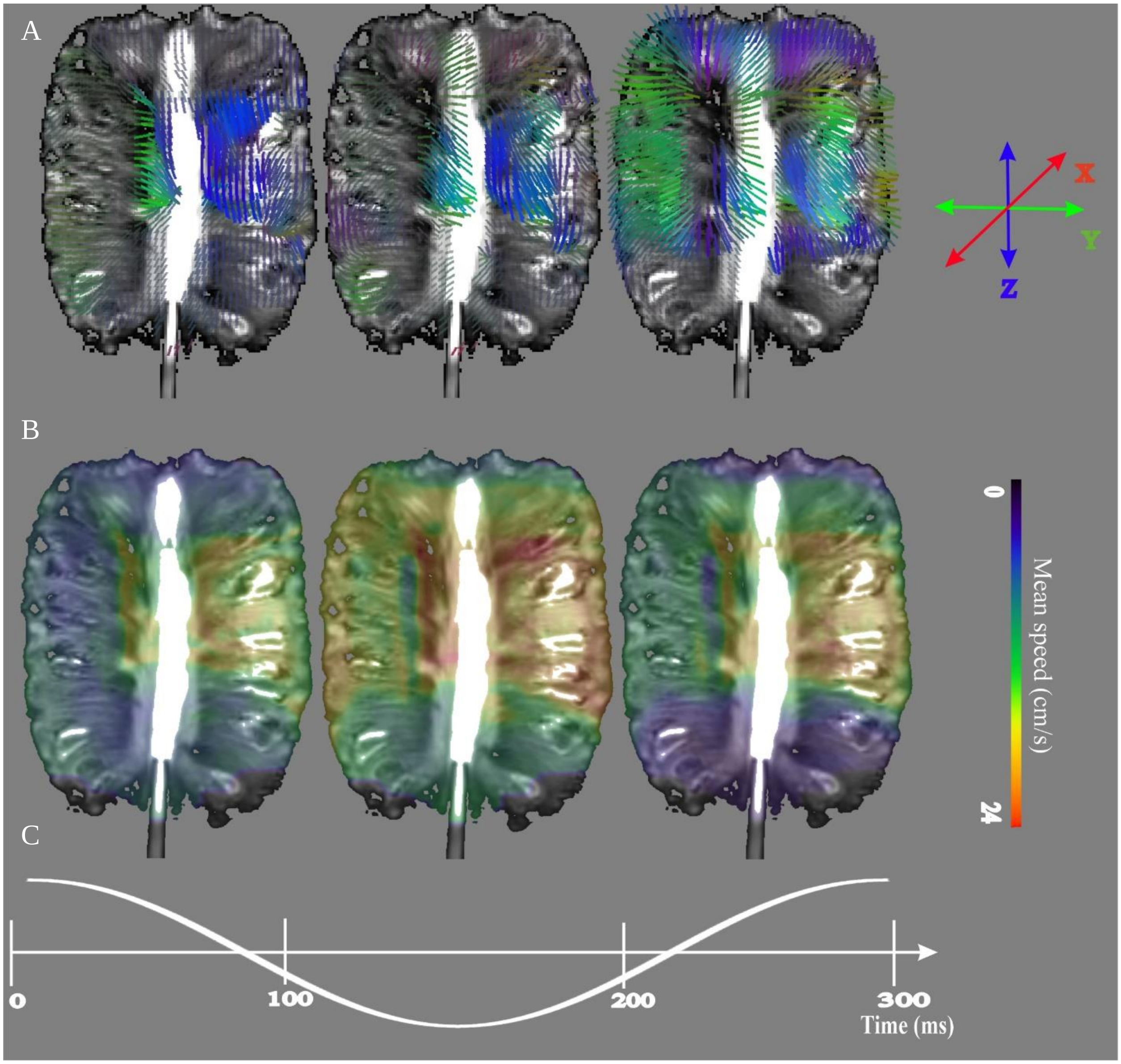


**Supplementary Figure 5. Mean velocity of optical flow analysis at stage 2 (20% maximal flow pump speed) through the pineapple.** (A) The 3D directional V maps of the stage 2 water flow impulse propagation are analyzed in three-time segments over the averaged 0.3 s cycle. (B) Speed magnitude $\mathbf{v}_s$ analysis of stage 2, over the three-time segments of the cycle. (C) Representative time domain signal of the stage 2 signal.

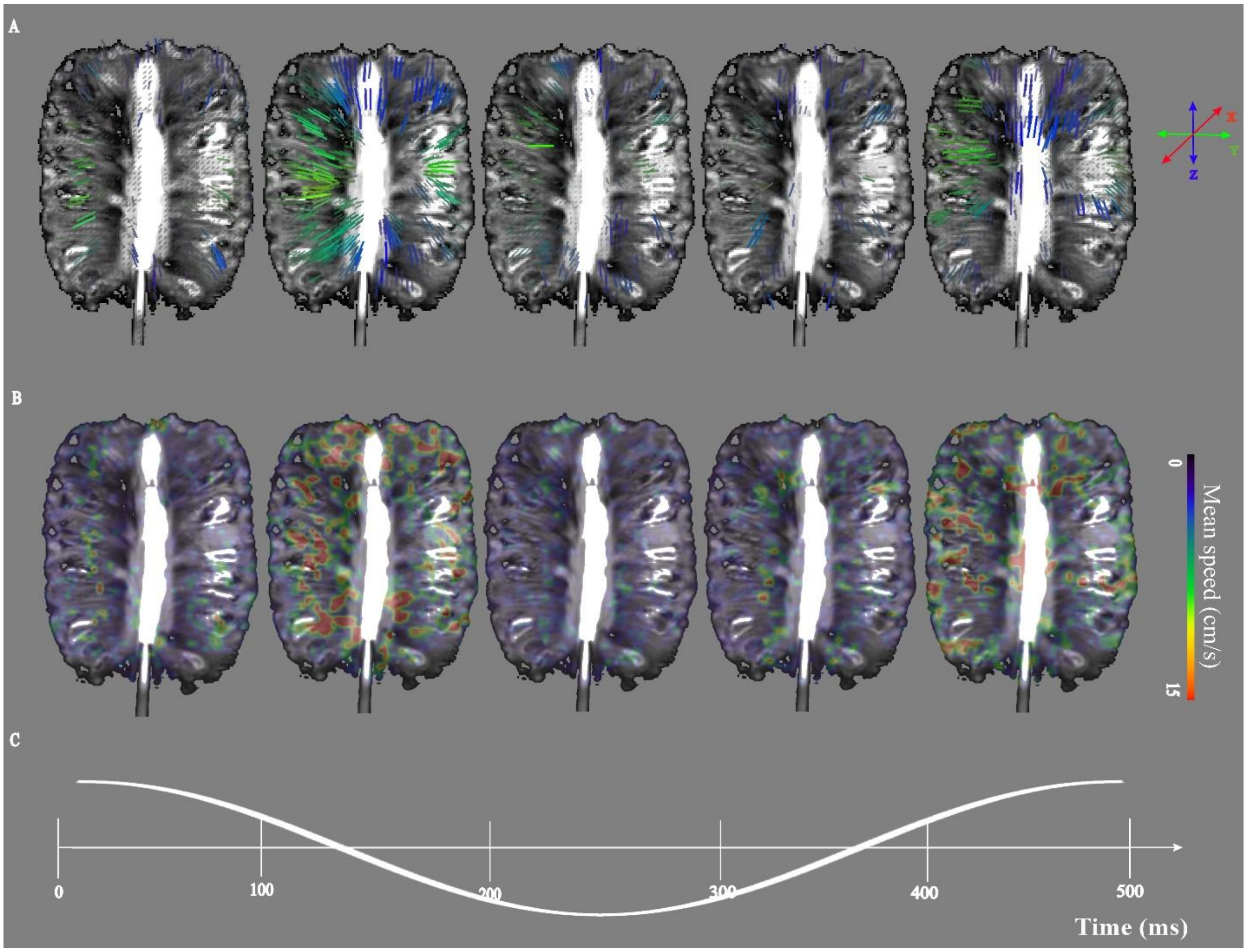


**Supplementary Figure 6. Mean velocity of sparse optical flow analysis at stage 1 (10% maximal flow pump speed) through the pineapple.** (A) The 3D directional V maps of the stage 1 water flow impulse propagation are analyzed in five-time segments over the averaged 0.5 s cycle. (B) Speed magnitude vs analysis of stage 1, over the five-time segments of the cycle. (C) Representative time domain signal of stage 1 signal.

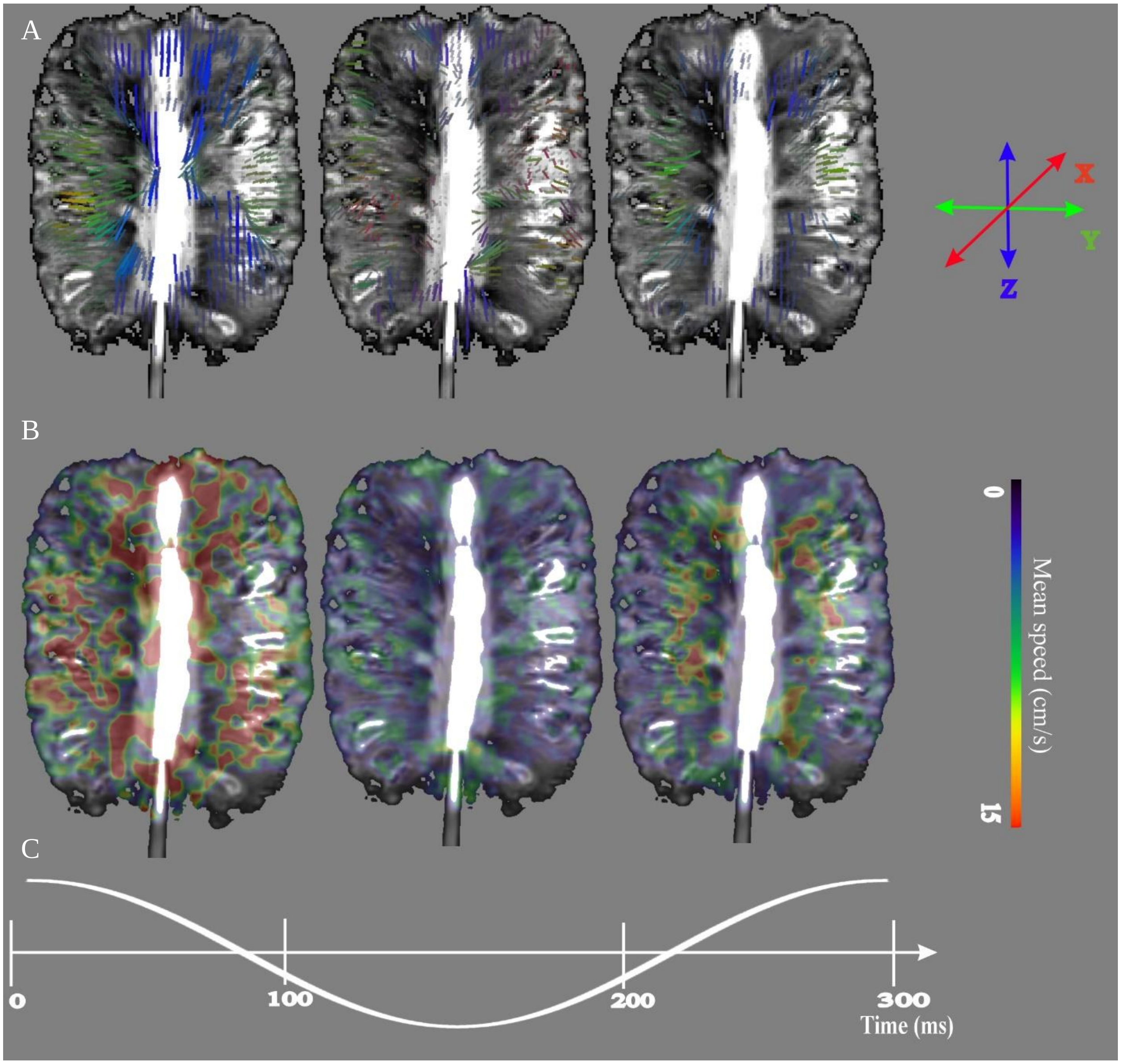


**Supplementary Figure 7. Mean velocity of optical flow analysis on stage 2 (20% maximal flow pump speed) through the pineapple.** (A) The 3D directional V maps of the stage 2 water flow impulse propagation are analyzed in three-time segments over the averaged 0.3 s cycle. (B) Speed magnitude $\mathbf{v_s}$ analysis of stage 2, over the three-time segments of the cycle. (C) Representative time domain signal of stage 2 signal.

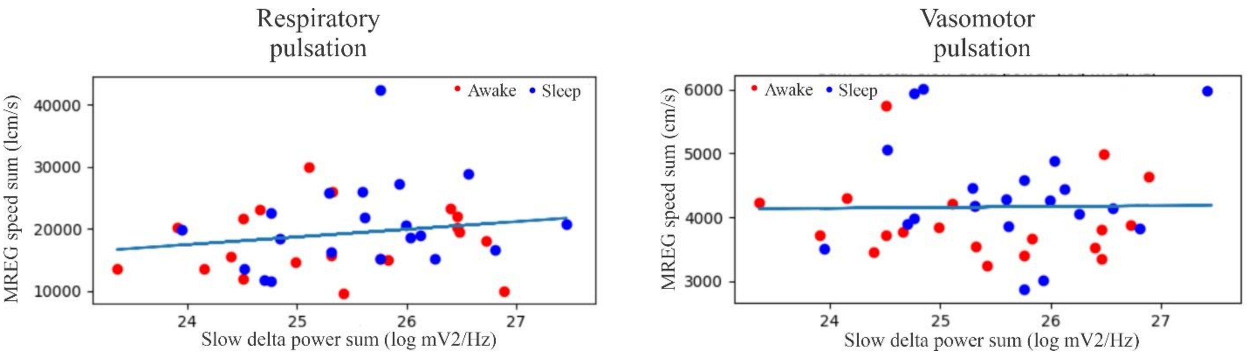


**Supplementary Figure 8. Speed and slow-delta power analysis of subjects in the two arousal states.** Mean MREG speed vs. slow Delta EEG power in each subject vs. mean slow-delta power of EEG data for respiratory and likewise vasomotor pulsations.
